# Quantitative assessment of multiple fish species around artificial reefs using environmental DNA metabarcoding

**DOI:** 10.1101/2021.05.05.442866

**Authors:** Masaaki Sato, Nariaki Inoue, Ryogen Nambu, Naoki Furuich, Tomohito Imaizumi, Masayuki Ushio

**Affiliations:** Fisheries Engineering Division, Fisheries Technology Institute, Japan Fisheries Research and Education Agency (FRA), Kamisu, Ibaraki, Japan; Fisheries Division, Japan International Research Center for Agricultural Sciences (JIRCAS), Tsukuba, Ibaraki, Japan; Hakubi Center, Kyoto University, Kyoto 606-8501 Japan; Center for Ecological Research, Kyoto University, Otsu, Shiga 520-2113 Japan

## Abstract

Since the early 1970s, many artificial reefs (ARs) have been deployed in Japanese coastal waters to create fisheries grounds. Recently, researchers began to use environmental DNA (eDNA) methods for biodiversity monitoring of aquatic species. A metabarcoding approach using internal standard DNAs (i.e., quantitative MiSeq sequencing) makes it possible to monitor eDNA concentrations of multiple species simultaneously. This method can improve the efficiency of monitoring AR effects on fishes. Our study investigated distributions of marine fishes at ARs and surrounding stations in the open oceanographic environment of Tateyama Bay, central Japan, using quantitative MiSeq sequencing and echo sounder survey. Using the quantitative metabarcoding method, we found higher quantities of fish eDNAs at the ARs than at surrounding stations and different fish species compositions between them. Comparisons with eco sounder survey also showed positive correlations between fish eDNA concentration and echo intensity, which indicates a highly localized signal of eDNA at each sampling station. These results suggest that quantitative MiSeq sequencing is a promising technique to complement conventional methods to monitor distributions of multiple fish species.

## Introduction

Artificial reefs (ARs) have been deployed worldwide to create fisheries grounds. Since the early 1970s, restrictions on overseas fishing have influenced high seas fisheries outside the exclusive economic zones (EEZ), and this situation led the Japanese government to implement a new fishery policy to develop and maintain coastal fisheries. In 1971, the Japanese government started the first national project of maintenance and development of coastal fisheries and has continued to implement similar projects, spending $ 5.4 billion to deploy ARs throughout coastal waters. Yamane^1^ indicated that the total number of Japanese AR projects and proportion of shallow fishing grounds with ARs in 1986 reached 6,443 and 9.3%, respectively. After establishing a large number of ARs in Japanese waters, underwater visual census, surveys using fishery gear, and echo sounder surveys have been used to examine fish aggregation and stock enhancement effects of ARs in Japanese waters^2–5^. These survey methods have advantages and disadvantages. An underwater visual census is effective for species identification, fish counts, and size estimation, but it requires identification skills in situ and/or a long investigation time for large areas. Surveys with bottom trawls and gillnets can obtain similar information on the fish to the underwater visual census, but these surveys exploit fish individuals. Echo sounders can survey large geographical areas quickly without fish collection, but with this method, it is difficult to identify fish species.

Recently, researchers have begun to use environmental DNA (eDNA), DNAs derived from environmental samples such as water, to investigate the distribution of aquatic organisms, including marine fishes^6,7^. In the case of macro-organisms, eDNA originates from metabolic waste, damaged tissue, dead individuals, or spawning events^8^, and the eDNA contains information about the species of the organisms that produced it^7^. This method does not need to exploit a body or body tissues of target species and can detect target organisms noninvasively. Also, this method requires less field time and is more likely to detect the presence of target species than the other survey methods^6,9^. Recently, the metabarcoding approach, using high-throughput sequencing technologies (e.g., Illumina MiSeq) and universal primer sets, has been utilized to reveal the fish diversity in a given area^6,10^. MiFish, a universal polymerase chain reaction (PCR) primer set, is especially popular for fish eDNA metabarcoding and has detected more fish species than other competing primers^11^.

Although the eDNA metabarcoding approach has become a cost- and labor-effective approach for estimating aquatic biodiversity, it has one limitation, i.e., the quantity of eDNA cannot be estimated from the number of eDNA sequence reads obtained by high-throughput sequencing partly due to several experimental problems such as PCR inhibitions^7^. However, some studies have proposed an approach using an internal standard DNA to overcome this limitation. Ushio et al.^7^ tested whether additions of internal standard DNAs (i.e., known amounts of short DNA fragments from fish species that have never been observed in a sampling area) to eDNA samples can create sample-specific regression lines between the numbers of sequence reads and DNA copies. Based on regression lines for each sample, they converted the sequence reads of DNAs from unknown fish species (i.e., non-standard fish species) to the number of fish DNA copies in each sample. This study showed that the estimated numbers of total and two dominant fish (Japanese anchovy and jack mackerel) DNA copies were significantly positively correlated with the numbers of DNA copies quantified by qPCR. Therefore, this method (i.e., quantitative MiSeq sequencing) can contribute to the quantitative assessment of multispecies fish eDNA. Some studies have shown that the eDNA quantity could be a proxy for the abundance or biomass of macro-organisms, not only in closed systems like tank and mesocosm^12^, but also in marine ecosystems^9,13^. The quantitative MiSeq sequencing with MiFish primer could be one of the most cost- and labor-effective approaches for the quantitative monitoring of multi species fish in marine and coastal ecosystems, including ARs.

Here, we investigated marine fish distributions around ARs in Tateyama Bay, central Japan, using eDNA metabarcoding, catch record of fishery, and eco sounder method. The quantitative MiSeq sequencing using internal standard DNAs and MiFish primers was applied to assess whether the distribution of multi fish species around ARs could be monitored based on the spatial variation in the estimated quantities of eDNA (i.e., identification of fish species and quantification of the number of fish eDNA copies simultaneously). Specifically, the following were tested: 1) whether fish communities detected by eDNA metabarcoding match with the catch record of the coastal fishery, 2) whether the DNA quantities of multiple fish species calculated by the quantitative MiSeq sequencing decrease with increasing distance from an AR, 3) whether the spatial variation of fish DNA quantities correlates to the relative distribution estimated by the acoustic survey.

## Materials and methods

### Study site, field survey, and in situ filtration

The field survey was performed in Tateyama Bay (34°60’N, 139°48’E), central Japan, in the proximity of the Kuroshio warm current facing the Pacific Ocean (Fig. 1). This area has many artificial reefs (ARs) created to improve fishing efficiency for fishers. Among the ARs, we focused on one high-rise steel AR (AR1), with a height of 30 m, where fish tended to aggregate (Fig. S1). Sampling stations were set up at the AR1 and at six linear distant points extending northeast and southwest. These stations were named E150, E500, E750, W150, W500, and W750, where “W” or “E” and the number of each station name represented northeast or southwest and distance in meters from the AR1, respectively (Table 1 and Fig. 1). Another station was set up at a second AR (AR2: 25 m height) 220 m from AR1 because we found AR2 by chance during the survey (Table 1 and Fig. 1), and it might affect the eDNA concentration at other stations.

**Figure 1.**
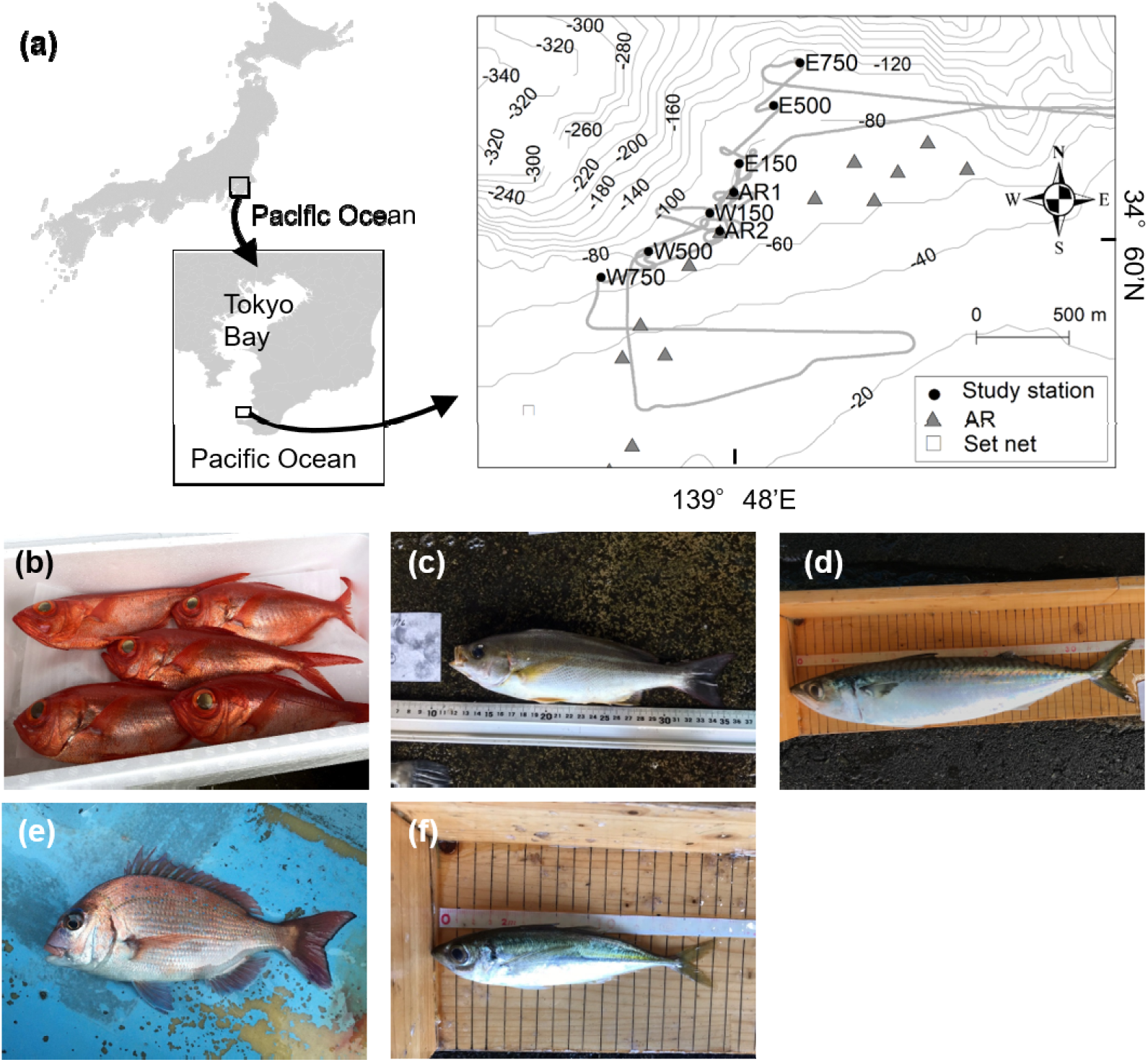
(a) Location of sampling stations, cruise track, and a set net in Tateyama Bay. Gray areas indicate landmasses, a gray bold line indicates cruise track, and gray thin lines indicate depth contours with an interval of 20 m. Pictures of the dominant species, (b) splendid alfonsino (*Beryx splendens*), (c) chicken grunt (*Parapristipoma trilineatum*), (d) chub mackerel (*Scomber japonicus*), (e) red seabream (*Pagrus major*), and (f) jack mackerel (*Trachurus japonicus*). Photograph credits: (b) Fumie Yamaguchi, (c, d, f) Yutaro Kawano, and (e) Masaaki Sato.

We conducted water sampling for eDNA analysis and performed an acoustic survey for estimating relative fish density using research vessel *Taka-maru* (Japan Fisheries Research and Education Agency: FRA) on May 23, 2018. We started the echo sounder survey at the southern part of the bay and continued it during the water sampling. Although the echo sounder survey could not differentiate between fish species, we collected this data to assess the association between the estimated concentration fish of eDNA and the echo intensity measured by the echo sounder. Water sampling began at E750, then continued along the transect line to E150, AR1, W150, W500, W750, before going back to AR2. At each sampling station, we collected 10 L of seawater from both the middle and bottom layers by one cast of two Niskin water samplers and measured vertical profiles of water temperature and salinity with a conductivity-temperature-depth sensor (RINKO profiler, JFE Advantech Co., Ltd.). Then, two 2L samples were subsampled from the seawater samples and immediately filtered using a combination of Sterivex filter cartridges (nominal pore size = 0.45 μm; Merck Millipore) through an aspirator (i.e., the two filters were subsets of a single water collection) in a laboratory on the research vessel. After filtration (approximately 15 min), an outlet port of the filter cartridge was sealed with an outlet luer cap, 1.5 ml RNAlater (ThermoFisher Scientific, Waltham, MA) was injected into the cartridge using a filtered pipette tip to prevent eDNA degradation, and an inlet port was also sealed with an inlet luer cap^14^. The Niskin water samplers were bleached before each water collection using a commercial bleach solution while filtering devices (i.e., filter funnels and measuring cups used for filtration) were bleached after every filtration. We filtered 2L MilliQ water with a filter funnel and measuring cup for every two or three sites as a field negative control to test for possible contamination. The filter cartridges were placed in a freezer immediately after filtration until eDNA extraction. Disposable latex or nitrile gloves were worn during sampling and replaced between each sampling station.

### DNA extraction and purification

Workspaces were sterilized prior to DNA extraction using 10% commercial bleach, and filter tip pipettes were used to safeguard against cross-contamination. Following the method developed by Miya et al.^14^, the eDNA was extracted and purified. Briefly, after removing RNAlater inside the cartridge using a centrifuge, proteinase-K solution was injected into the cartridge from the inlet port, and the port was re-capped with the inlet lure cap. The eDNA captured on the filter membrane was extracted by constant stirring of the cartridge at a speed of 20 rpm using a roller shaker placed in an incubator heated at 56°C for 20 min. The eDNA extracts were transferred to a 2-ml tube from the inlet of the filter cartridges by centrifugation. The collected DNA was purified using a DNeasy Blood & Tissue Kit (Qiagen) following the manufacturer’s protocol. After the purification, DNA was eluted using 100 μl of the elution buffer (buffer AE). All DNA extracts were frozen at −20°C until paired-end library preparation.

### Preparation of internal standard DNAs

Five artificially designed and synthetic internal standard DNAs, which were similar but not identical to the region of any existing fish mitochondrial 12S rRNA, were included in the library preparation process to estimate the number of fish DNA copies (i.e., quantitative MiSeq sequencing)^7,15^. They were designed to have the MiFish primer□binding regions as those of known existing fishes and to have the conserved regions in the insert region. Variable regions in the insert region were replaced with random bases so that no known existing fish sequences had the same sequences as the standard sequences. The standard DNA size distribution of the library was estimated using an Agilent 2100 BioAnalyzer (Agilent, Santa Clara, CA, USA), and the concentration of double-stranded DNA of the library was quantified using a Qubit dsDNA HS assay kit and a Qubit fluorometer (ThermoFisher Scientific, Waltham, MA, USA). Based on the quantification values obtained using the Qubit fluorometer, the copy number of the standard DNAs was adjusted as follows: Std. A (100 copies/μl), Std. B (50 copies/μl), Std. C (25 copies/μl), Std. D (12.5 copies/μl) and Std. E (2.5 copies/μl). Then, these standard DNAs were mixed.

### Paired-end library preparation

Four PCR - level negative controls (i.e., each two with and without internal standard DNAs) were employed for MiSeq run to monitor contamination during the experiments. The first-round PCR (1st PCR) was carried out with a 12-μl reaction volume containing 6.0 μl of 2 × KAPA HiFi HotStart ReadyMix (KAPA Biosystems, Wilmington, WA, USA), 0.7 μl of each primer (5 μM), 2.6 μl of sterilized distilled H_2_O, 1.0 μl of standard DNA mix and 1.0 μl of template. Note that the standard DNA mix was included for each sample. The final concentration of each primer was 0.3 μM. We used a mixture of the following four PCR primers modified from original MiFish primers^16^: MiFish-U-forward (5’-ACA CTC TTT CCC TAC ACG ACG CTC TTC CGA TCT NNN NNG TCG GTA AAA CTC GTG CCA GC) and MiFish-U-reverse (5’-GTG ACT GGA GTT CAG ACG TGT GCT CTT CCG ATC TNN NNN CAT AGT GGG GTA TCT AAT CCC AGT TTG-3’), MiFish-E-forward (5’-ACA CTC TTT CCC TAC ACG ACG CTC TTC CGA TCT NNN NNG TTG GTA AAT CTC GTG CCA GC-3’), and MiFish-E-reverse (5’-GTG ACT GGA GTT CAG ACG TGT GCT CTT CCG ATC TNN NNN CAT AGT GGG GTA TCT AAT CCT AGT TTG-3’). These primer pairs co-amplify a hypervariable region of the fish mitochondrial 12S rRNA gene (around 172 bp; hereafter called MiFish sequence) and append primer-binding sites (5’ ends of the sequences before six Ns) for sequencing at both ends of the amplicon. The five random bases were used to enhance cluster separation on the flow cells during initial base call calibrations on the MiSeq platform. The thermal cycle profile after an initial 3 min denaturation at 95°C was as follows (35 cycles): denaturation at 98°C for 20 s; annealing at 65°C for 15 s; and extension at 72°C for 15 s, with a final extension at the same temperature for 5 min. Eight replications were performed for the 1st PCR, and the replicates were pooled to minimize the PCR dropouts. The 1st PCR products from the eight tubes were pooled in a single 1.5-ml tube. Then, we sent the 1st PCR products to IDEA consultants, Inc. to outsource the following MiSeq sequencing processes. The pooled products were purified and size-selected for 200-400bp using a SPRIselect (Beckman Coulter inc.) to remove dimers and monomers following the manufacturer’s protocol.

The second-round PCR (2nd PCR) was carried out with a 24 μl reaction volume containing 12 μl of 2 × KAPA HiFi HotStart ReadyMix, 2.8 μl of each primer (5 μM), 4.4 μl of sterilized distilled H2O, and 2.0 μl of template. We used the following two primers to append the dual-index sequences (8 nucleotides indicated by Xs) and flowcell-binding sites for the MiSeq platform (5’ ends of the sequences before eight Xs): 2nd-PCR-forward (5’-AAT GAT ACG GCG ACC ACC GAG ATC TAC ACX XXX XXX XAC ACT CTT TCC CTA CAC GAC GCT CTT CCG ATC T-3’); and 2nd-PCR-reverse (5’-CAA GCA GAA GAC GGC ATA CGA GAT XXX XXX XXG TGA CTG GAG TTC AGA CGT GTG CTC TTC CGA TCT-3’). The thermal cycle profile after an initial 3 min denaturation at 95°C was as follows (12 cycles): denaturation at 98°C for 20 s; combined annealing and extension at 72°C for 15 s, with a final extension at 72°C for 5 min. The concentration of each second PCR product was measured by quantitative PCR using TB Green Fast qPCR Mix (Takara inc.). Each sample was diluted to a fixed concentration and combined (i.e., one pooled 2nd PCR product that included all samples). The pooled 2nd PCR product was size-selected to approximately 370 bp using BluePippin (SAGE SCIENCE). The size-selected library was purified using the Agencourt AMPure XP beads, adjusted to 4 nM by quantitative PCR using TB Green Fast qPCR Mix (Takara inc.), and sequenced on the MiSeq platform using a MiSeq v2 Reagent Kit (2 × 150 bp).

### Data preprocessing and taxonomic assignment

The raw MiSeq data were converted into FASTQ files using the bcl2fastq program provided by Illumina (bcl2fastq v2.18). The FASTQ files were then demultiplexed using the command implemented in Claident^17^. We adopted this process rather than using FASTQ files demultiplexed by the Illumina MiSeq default program in order to remove sequences whose 8□mer index positions included nucleotides with low-quality scores (i.e., Q-score < 30).

The processed reads were subjected to a BLASTN search against the full NCBI database. If the sequence similarity between queries and the top BLASTN hit was < 98.5% and the sequence length was less than ≤ 150 bp, the unique sequence was not subjected to the following analyses. After BLASTN searches, assembled sequences assigned to the same species were clustered, and we considered the clustered sequences as operational taxonomic units (OTUs). To remove possible contaminants, we removed the OTUs whose sequence reads were < 0.05 % of the total reads as a noise for each sample.

### Catch record of coastal fishery

To confirm the feasibility of MiFish metabarcoding, we compared the fish community of MiFish metabarcoding with a catch record of a net set near the study site (Fig.1). The detection rate was defined as the proportion of species detected by metabarcoding compared to the species collected by the net. Because our eDNA survey was conducted in May 2018, we used catch-record data from the same period (May 2018).

### Estimation of DNA copy numbers

According to Ushio et al.^7^ and Ushio^15^, the procedure to estimate DNA copy numbers consisted of two parts: (i) linear regression analysis to examine the relationship between sequence reads and the copy numbers of the internal standard DNAs for each sample and (ii) the conversion of sequence reads of non-standard fish DNAs to estimate the copy numbers using the result of the linear regression for each sample.

Linear regressions were used to examine how many sequence reads were generated from one fish DNA copy through the library preparation process. Note that a linear regression analysis between sequence reads and standard DNAs was performed for each sample, and the intercept was set as zero. The regression equation was: MiSeq sequence reads = regression slope × the number of standard DNA copies [/μl]. The number of linear regressions performed was thirty-three (=the number of fish DNA samples + field negative controls with standard DNAs), and thus thirty-three regression slopes were estimated in total.

The sequence reads of non-standard fish DNAs were converted to copy numbers using sample-specific regression slopes estimated by the above regression analysis. The number of non-standard fish DNA copies was estimated by dividing the number of MiSeq sequence reads by a sample-specific regression slope (i.e., the number of DNA copies = MiSeq sequence reads/regression slope). A previous study demonstrated that these procedures provide a reasonable estimation of DNA copy numbers using high-throughput sequencing^7^.

### Acoustic survey of fish school using echo sounder

We estimated relative fish density using a quantitative echo sounder following the acoustic survey of Yamamoto et al.^9^. We used the echo sounder, KFC-6000 (Sonic Co. Tokyo, Japan), with a transducer (frequency, 38 kHz; beam type, split-beam; beam width, approximately 7°; pulse duration, 0.3 ms; ping rate, 0.4 s). The transducer was set at the bottom of the research vessel at a depth of 2.1 m to avoid cavitation bubbles generated by the research vessel. The acoustic devices were operated during the entire survey cruise, and all signals were recorded. The average ship speed was ~10 knots between sampling stations, although the ship slowed down when approaching a sampling station and completely stopped when water samples were being collected.

We eliminated noise from the obtained echo intensity data using Echoview ver. 11.0 (Echo-view Software Pty. Ltd., Tasmania, Australia). Signals between the sea bottom and 1.0 m above the sea bottom were eliminated to avoid possible confounding with the acoustic dead zone. In addition, the signas of Niskin water samplers and RINKO profiler were also eliminated. Finally, we eliminated the signals of bubbles generated by the movement of the screw propeller. After eliminating these noises, we re-obtained echo intensity data to assess the association with eDNA concentration. Note that the obtained echo intensity data would include echo signals from a variety of fish species.

### Data analyses

We performed three types of univariate analysis using generalized linear models (GLMs). First, we used likelihood ratio tests to assess variation in the eDNA copy numbers of total fish, demersal fish, pelagic fish, and five dominant OTUs, the number of OTUs, as well as echo intensity among sampling stations and between vertical positions of water sampling. When the effects of sampling stations were significant, the Tukey-HSD test was used to determine the significant differences between sampling stations. The second analysis assessed the number of OTUs and eDNA copy numbers of total, demersal, pelagic, and five dominant OTUs in relation to distance from the AR1, vertical position of water samples (i.e., middle or bottom layers, represented as ‘depth’), and the interaction between the distance and vertical position. Although our sampling stations contained two ARs, we focused on AR1 in the second analysis because more fish aggregated around AR1 and transport of eDNAs from AR1 may have more impact than those from AR2. The third analysis examined the relationships between eDNA copy numbers (each eDNA copy number of total fish, demersal fish, pelagic fish, or five dominant OTUs described below) and echo intensity (S_V_). Because echo intensity can correlate to distance from the AR1, the second and third analyses were separately performed. Based on Fishbase^18^, each fish OTU was classified into demersal and pelagic species. For the analyses, we used the Poisson distribution with a log-link function for OTU number because it was count data, while a gamma distribution with a log-link was used for eDNA copy numbers of total fish, pelagic fish, demersal fish, and *Scomber* spp. (one dominant OTU) because these values are continuous and greater than zero. A mixture of binomial and gamma distributions, which is often called a “gamma hurdle model,” was also used as in previous studies^19,20^ for eDNA copy numbers of remaining dominant OTUs because these values are continuous and contain zero values. Bayesian information criterion (BIC) was used to select the correct model among candidate models for the second and third analyses^21^.

We also performed multivariate analyses to determine differences in the fish community structures between sampling stations/vertical positions. Community structures of each sampling were visualized using non-metric dimensional scaling (NMDS) based on the Bray–Curtis dissimilarity index of the copy numbers of each OTUs. Similarity tests between two variables (sampling stations and vertical positions) were conducted using nonparametric multivariate analysis of variance (NPMANOVA). All statistical analyses were performed using RStudio version 1.1.423 (R version 3.6.1) with several packages. The packages used were: (a) glmmTMB^22^ for regression models; (b) multcomp^23^ for Tukey-HSD test; (c) MuMIn^24^ for BIC model selection, (d) vegan^25^ for multivariate analysis; (e) ggplot2^26^ and (f) ggpubr^27^ for figure creation.

## Results

### Sequence reads

The MiSeq paired□end sequencing (2 × 150 bp) of the 34 libraries for this study (containing 32 PCR replicates, one negative field control, and one PCR-level control), together with 142 libraries from another study (total number of libraries = 176), yielded a total of 15.98 million reads, with 95.3% base calls containing Phred quality scores of ≥ 30.0 (Q30; error rate = 0.1% or base call accuracy = 99.9%).

For this study, we obtained 4,862,616 MiSeq reads, of which 3,528,543 were high-quality, merged reads. Number of non-standard fish Miseq reads were 2,153,777 out of the 3,528,543 (61.0%). In the MiSeq run, 18.8%–98.6% of sequence reads were from non-standard fish DNAs of field samples, and 0% was from those of a field negative control. The final list included 95 OTUs distributed across 49 families and 83 genera (Table S1).

### Comparison of eDNA metabarcoding with fish catch record

The fish catch record of a set net indicated that 289 thousand kg of total fish biomass belonging to 41 classifications (39 species and two classifications of closely related species) was caught in May 2018. The catch biomass was highly variable between the classifications. For example, 255 thousands kg of *Scomber japonicus* was recorded, whereas *Fistularia petimba* was 0.2 kg only (Table S2). Of the 41 classifications caught by the set net, eDNA metabarcoding detected 25 species. Three classifications of the 41 classifications were not recorded at the species level in the list of the fish catch of the set net, while the metabarcoding cannot assign OTUs at the species level for the *Scomber japonicus* and *S. australasicus* because they are so closely related. Thus, excluding these five species, our metabarcoding eventually detected 58.3% (= 21/36) of the fishes caught by the set net. Among the five dominant OTUs detected by MiFish metabarcoding described below (*B. splendens*, *P. trilineatum*, *Scomber* spp., *P. major*, and *T. japonicus*), four OTUs except *B. splendens* appeared in the list of the fish catch.

### Relationship between the copy numbers and sequence reads of the standard DNA

The relationships between the sequence reads and copy numbers of standard DNAs were examined by linear regression analysis (Fig. 2). Within each sample, the sequence reads linearly and positively correlated with the copy numbers of standard DNAs (Fig. 2a). R^2^ values of the regression lines ranged from 0.71 to 0.99, and more than 75% of the samples had values over 0.9 (Fig. 2b). These results suggested that the number of sequence reads was generally proportional to the number of copies of DNA initially added to the first PCR reaction within a single sample. The slopes of the linear regressions varied depending on the sample, suggesting that the amplification efficiency or removal amounts of DNAs during the purification of PCR products were dependent on the individual sample. These sample-specific slopes were used to convert sequence reads to the copy numbers of fish OTUs. Based on the estimated copy numbers of each OTU, five dominant OTUs were *Beryx splendens*, *Parapristipoma trilineatum*, *Scomber* spp. (*S. japonicus* or *S. australasicus*), *Pargus major*, and *Trachurus japonicus* (Fig. 1b–f).

**Figure 2.**
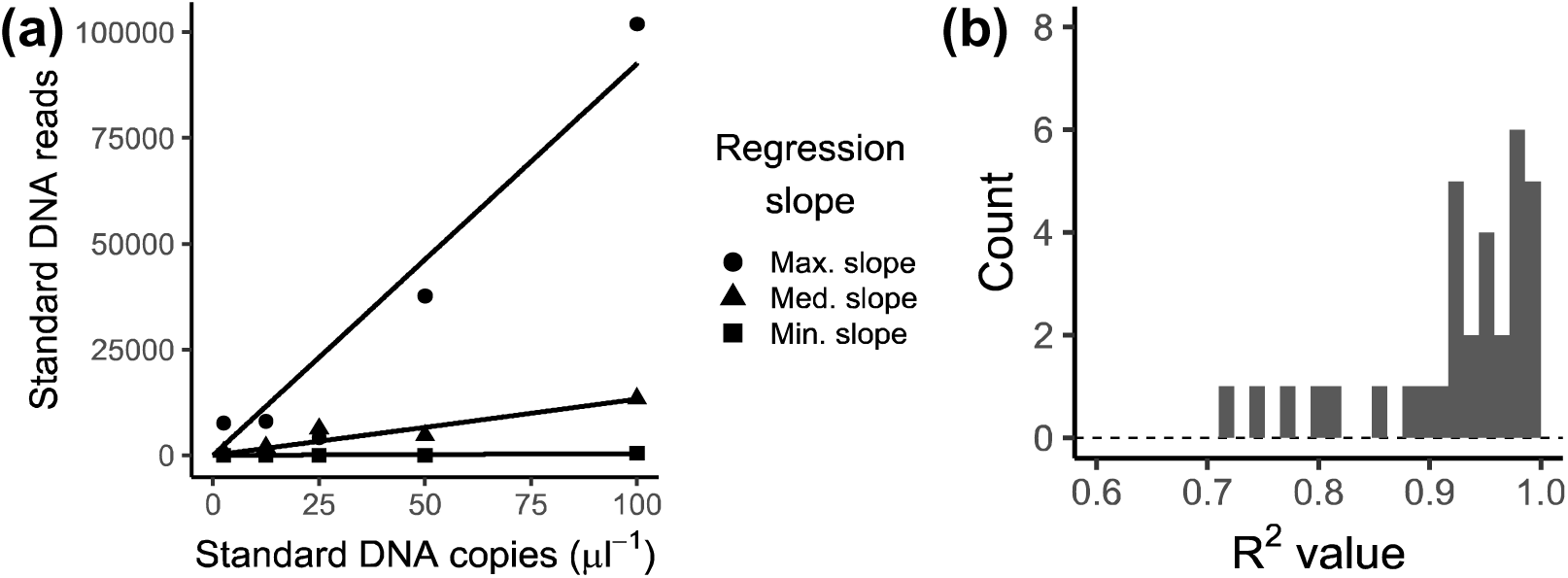
The relationships between sequence reads and the copy numbers of standard DNAs according to Ushio et al.^7^. (a) The relationships between sequence reads and the copy numbers of standard DNAs with the maximum, median, and minimum slopes. (b) Distribution of R^2^ values of the regression lines.

Levels of contaminations were assessed using the field negative control. DNA copy numbers detected in the field negative control (n = 1) were 0% of mean DNA copy numbers of field-positive samples, suggesting that there was no amount of contamination during the library preparation process.

### Site and vertical variation in fish eDNA quantities, the number of OTUs, and echo intensity

The estimated copy numbers of fish OTUs were summed to estimate the total, demersal, and pelagic fish eDNA quantities. The average eDNA copy numbers of total fish, demersal fish, and pelagic fish were 64.3, 44.6, and 19.7 copies/ml water, respectively, while those of five dominant OTUs, *B. splendens*, *P. trilineatum*, *Scomber* spp., *P. major*, and *T. japonicus*, were 20.3, 13.8, 12.9, 5.5, and 3.2 copies/ml water, respectively. Likelihood ratio tests showed that the eDNA copy numbers significantly varied among sampling stations for total, demersal, pelagic fishes, *P. trilineatum*, *Scomber* spp., *P. major*, and *T. japonicus*, whereas no significant variation was detected for *B. splendens* (Table S3). Tukey-HSD tests indicated that significantly higher total and demersal fish eDNA copy numbers were observed at AR1 than at W150–750 and E 500–750, while the pelagic fish eDNA copy number was higher at AR1 than at E750 (Fig. S1). Also, the total, demersal, and pelagic fish eDNA quantities were significantly higher at AR2 than at E750. For dominant fish OTUs, *P. trilineatum*, *P. major*, and *T. japonicus* eDNA copies were significantly higher at AR1 than at other stations, while *Scomber* spp. eDNA quantities were higher at AR1 and AR2 than at E750. Echo intensity (S_V_) ranged from 4.53×10^-8^ to 1.38×10^-6^, and a likelihood ratio test indicated significant variation in echo intensity among the stations. Like the fish eDNA quantities, the echo intensity was higher at AR1 than at the surrounding stations. Meanwhile, the number of OTUs detected at each station ranged from 9 to 27, with a mean of 16 OTUs, and there was no significant variation among the stations (Table S3).

Aside from echo intensity, likelihood ratio tests indicated that all the types of fish eDNA quantities and number of OTUs were significantly higher in the bottom samples than in the middle ones (Table S3 and Fig. S2).

### Effects of distance from an artificial reef on fish eDNA quantities and the number of OTUs

Distance from the AR1 and vertical position were included in the Gamma component of the minimum BIC model for total fish, demersal fish, pelagic fish, and *Scomber* spp. eDNA quantities (Table 2). Estimated parameters indicated that these eDNA quantities decreased with increasing distance from the AR1 and the eDNA quantities were lower in the middle samples than in the bottom ones. For *Parapristipoma trilineatum*, *Pargus major*, and *Trachurus japonicus* eDNA quantities, distance from the AR1, vertical position, and their interaction were included in the Gamma component of the minimum BIC model. Estimated parameters indicated that these eDNA quantities decreased with increasing distance from AR1, but the intercepts and slopes of the relationship with distance differed between the layers. The eDNA quantities of *P. major* were lower in the middle layer at 0 m from AR1, while those of *P. trilineatum* and *T. japonicus* were higher in the middle samples at 0 m from AR1 (Fig. 3). Slopes for *P. trilineatum*, and *T. japonicus* eDNA quantities were steeper in the middle layer, while that for *P. major* eDNA was steeper in the bottom layer. Only the vertical position was included in the Gamma component of the minimum BIC model for the number of OTUs and *Beryx splendens* eDNA quantity, and these values were higher in the bottom layer than in the middle layer (Table 2 and Fig. 3).

**Figure 3.**
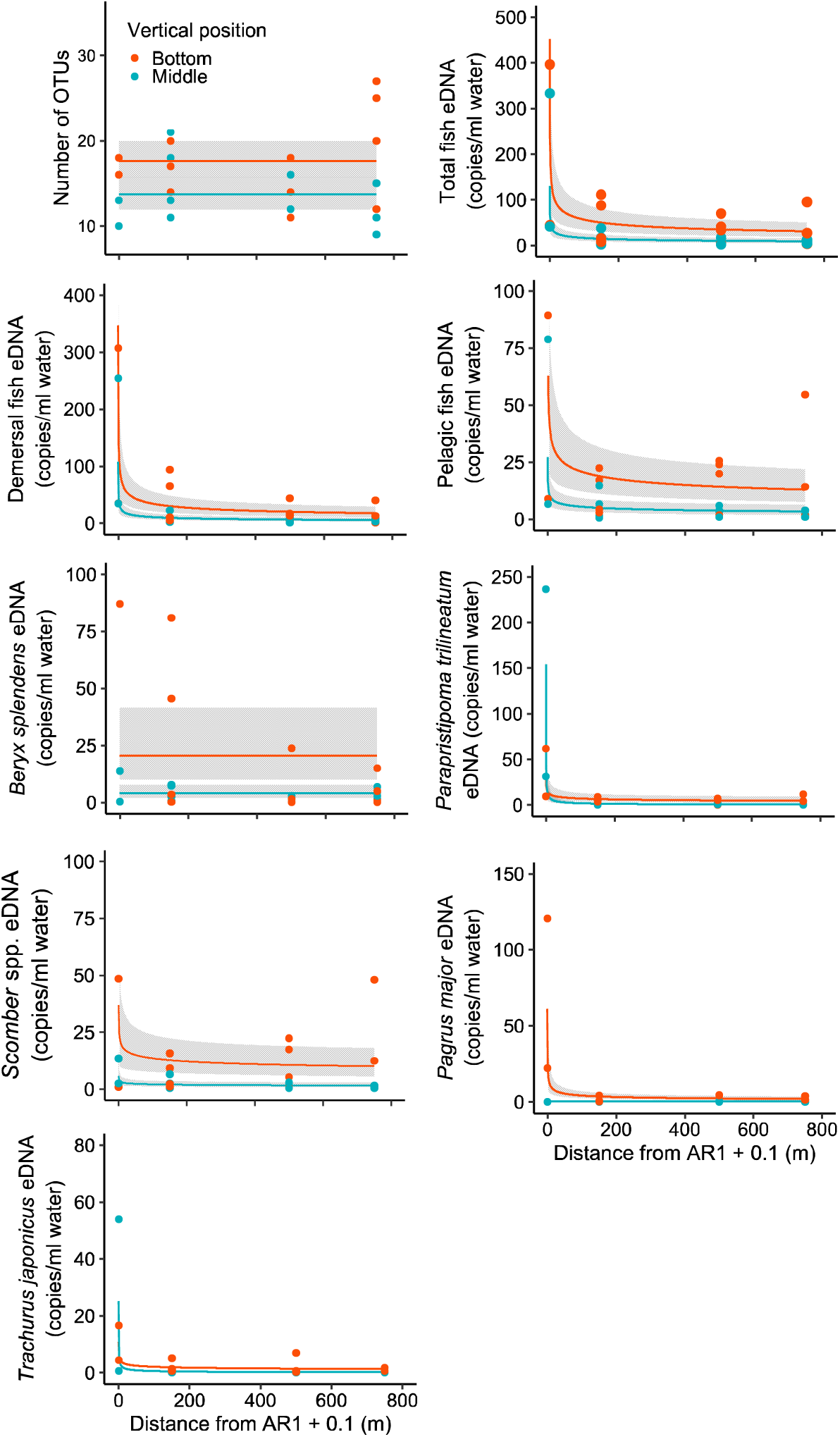
The relationships between distance from AR1 as well as number of OTUs and eDNA copies of total fish, demersal fish, pelagic fish, *Beryx splendens*, *Parapristipoma trilineatum*, *Scomber* spp., *Pagrus major*, and *Trachurus japonicus*. The black line and the gray shaded area represent a median value and 95% confidence interval, respectively, as determined from the generalized linear model. Red and turquoise blue circles indicate values in the bottom and middle sampling layers.

**Figure 4.**
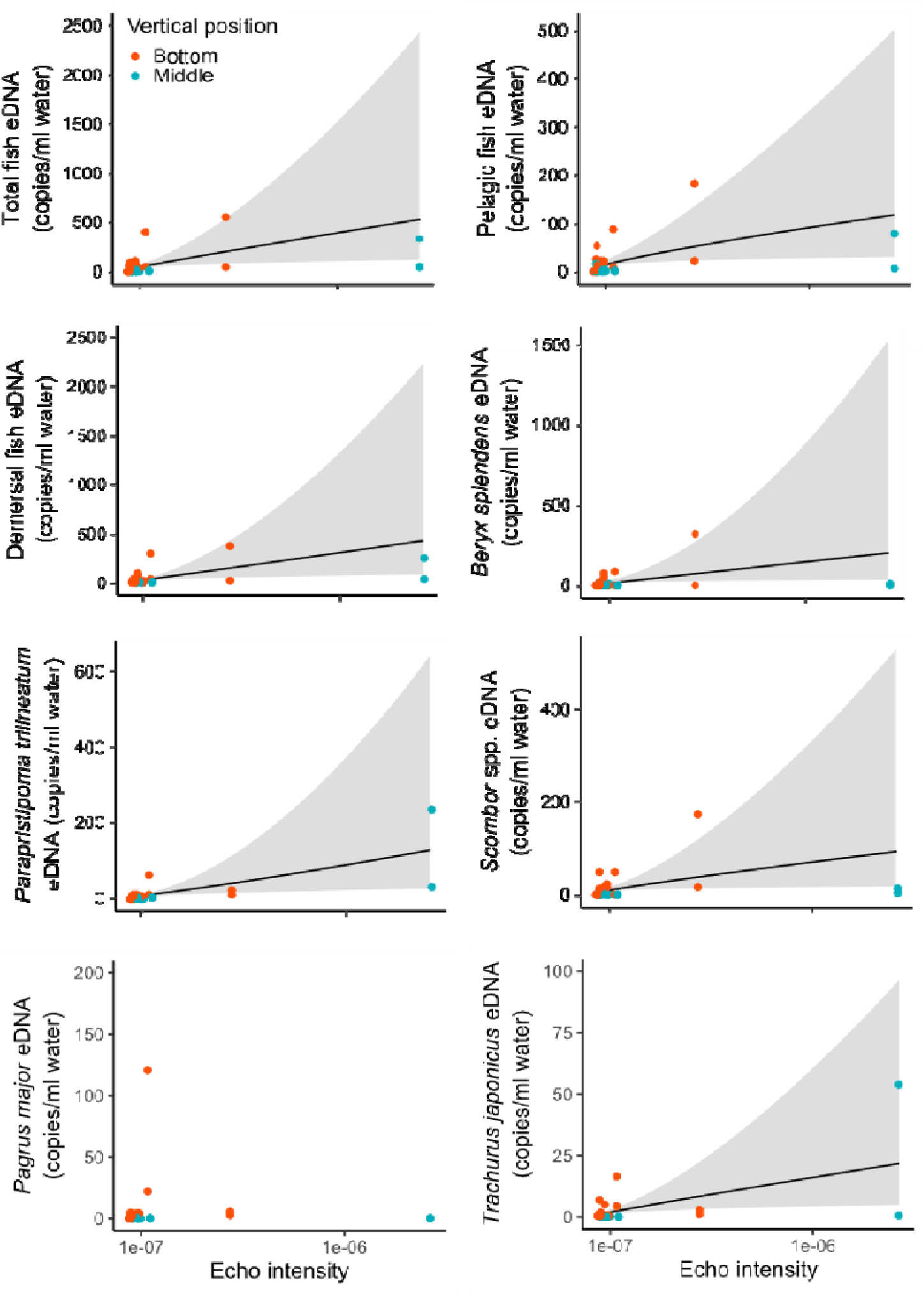
The relationships between eco intensity and eDNA copies of total fish, demersal fish, pelagic fish, *Beryx splendens*, *Parapristipoma trilineatum*, *Scomber* spp., *Pagrus major*, and *Trachurus japonicus*. The black line and the gray shaded area represent a median value and the 95% confidence interval, respectively, as determined from the generalized linear model. Red and turquoise blue circles indicate values in the bottom and middle sampling layers.

**Figure 5.**
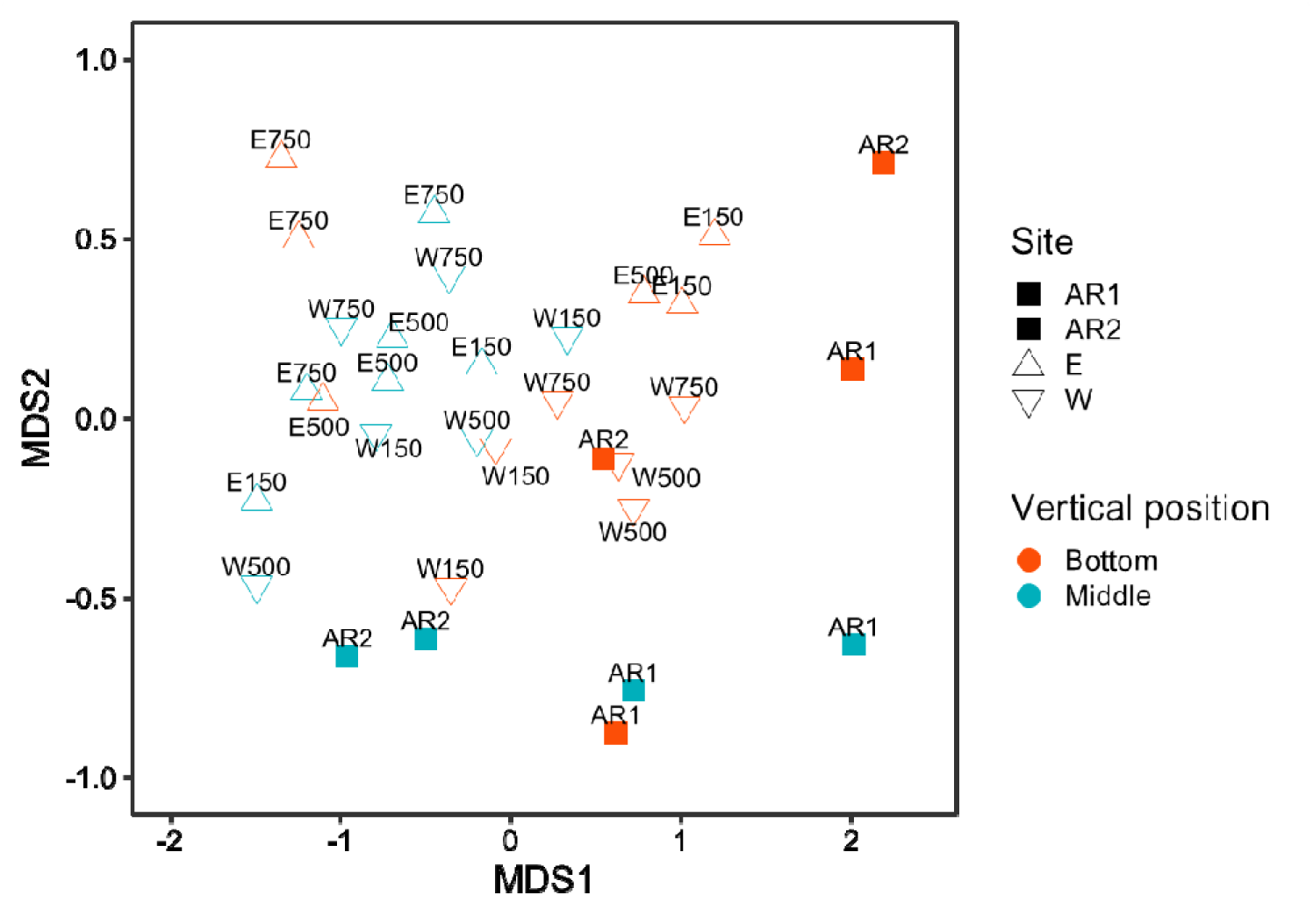
Nonmetric multidimensional scaling (NMDS) for fish communities detected at sampling stations. NMDS stress was 0.096.

**Table 1.**
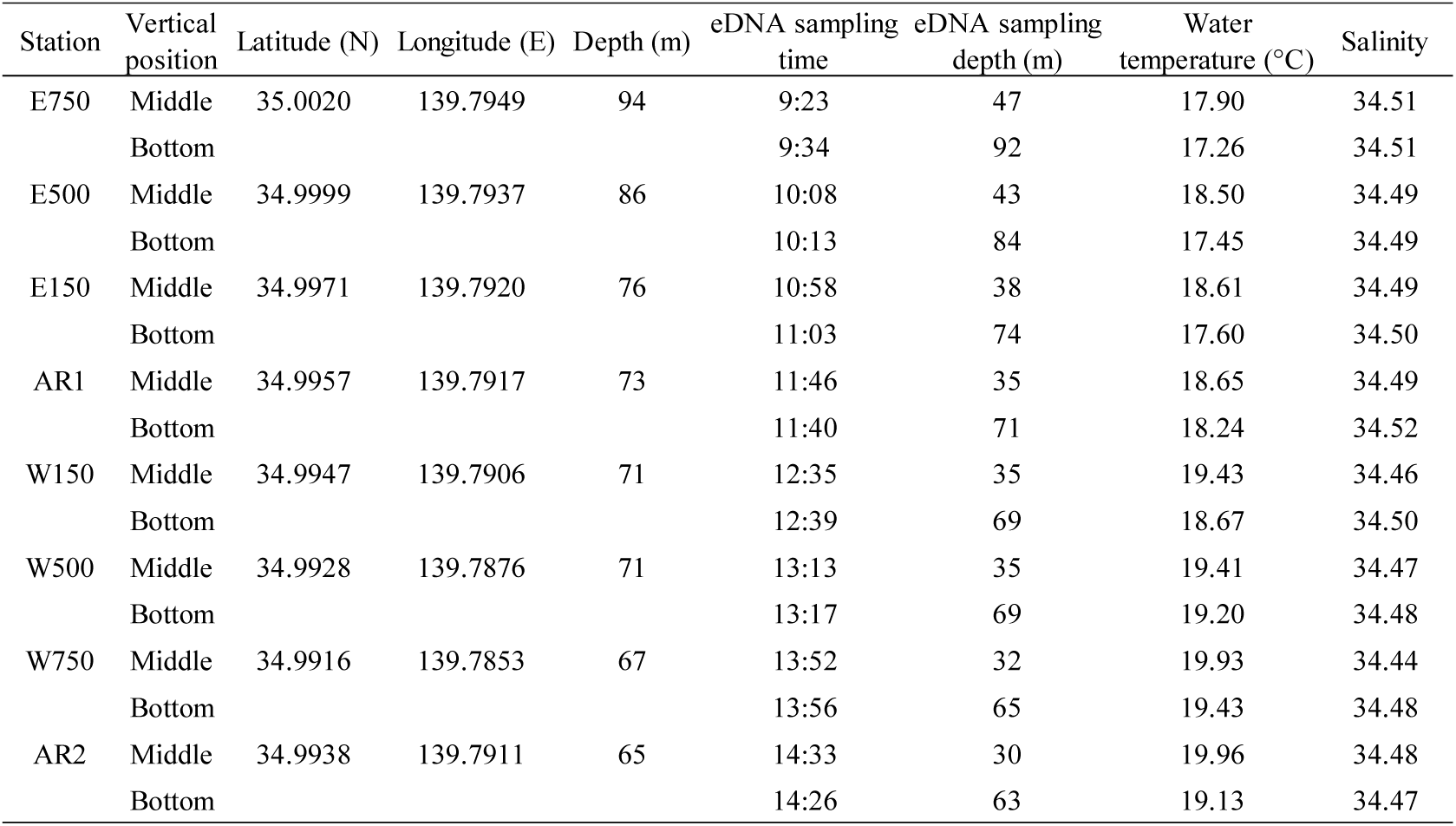
Details of the eight sampling stations for fish eDNA survey on May 23, 2018

| Station | Vertical position | Latitude (N) | Longitude (E) | Depth (m) | eDNA sampling time | eDNA sampling depth (m) | Water temperature (°C) | Salinity |
| --- | --- | --- | --- | --- | --- | --- | --- | --- |
| E750 | Middle | 35.0020 | 139.7949 | 94 | 9:23 | 47 | 17.90 | 34.51 |
|  | Bottom |  |  |  | 9:34 | 92 | 17.26 | 34.51 |
| E500 | Middle | 34.9999 | 139.7937 | 86 | 10:08 | 43 | 18.50 | 34.49 |
|  | Bottom |  |  |  | 10:13 | 84 | 17.45 | 34.49 |
| E150 | Middle | 34.9971 | 139.7920 | 76 | 10:58 | 38 | 18.61 | 34.49 |
|  | Bottom |  |  |  | 11:03 | 74 | 17.60 | 34.50 |
| AR1 | Middle | 34.9957 | 139.7917 | 73 | 11:46 | 35 | 18.65 | 34.49 |
|  | Bottom |  |  |  | 11:40 | 71 | 18.24 | 34.52 |
| W150 | Middle | 34.9947 | 139.7906 | 71 | 12:35 | 35 | 19.43 | 34.46 |
|  | Bottom |  |  |  | 12:39 | 69 | 18.67 | 34.50 |
| W500 | Middle | 34.9928 | 139.7876 | 71 | 13:13 | 35 | 19.41 | 34.47 |
|  | Bottom |  |  |  | 13:17 | 69 | 19.20 | 34.48 |
| W750 | Middle | 34.9916 | 139.7853 | 67 | 13:52 | 32 | 19.93 | 34.44 |
|  | Bottom |  |  |  | 13:56 | 65 | 19.43 | 34.48 |
| AR2 | Middle | 34.9938 | 139.7911 | 65 | 14:33 | 30 | 19.96 | 34.48 |
|  | Bottom |  |  |  | 14:26 | 63 | 19.13 | 34.47 |

**Table 2.**
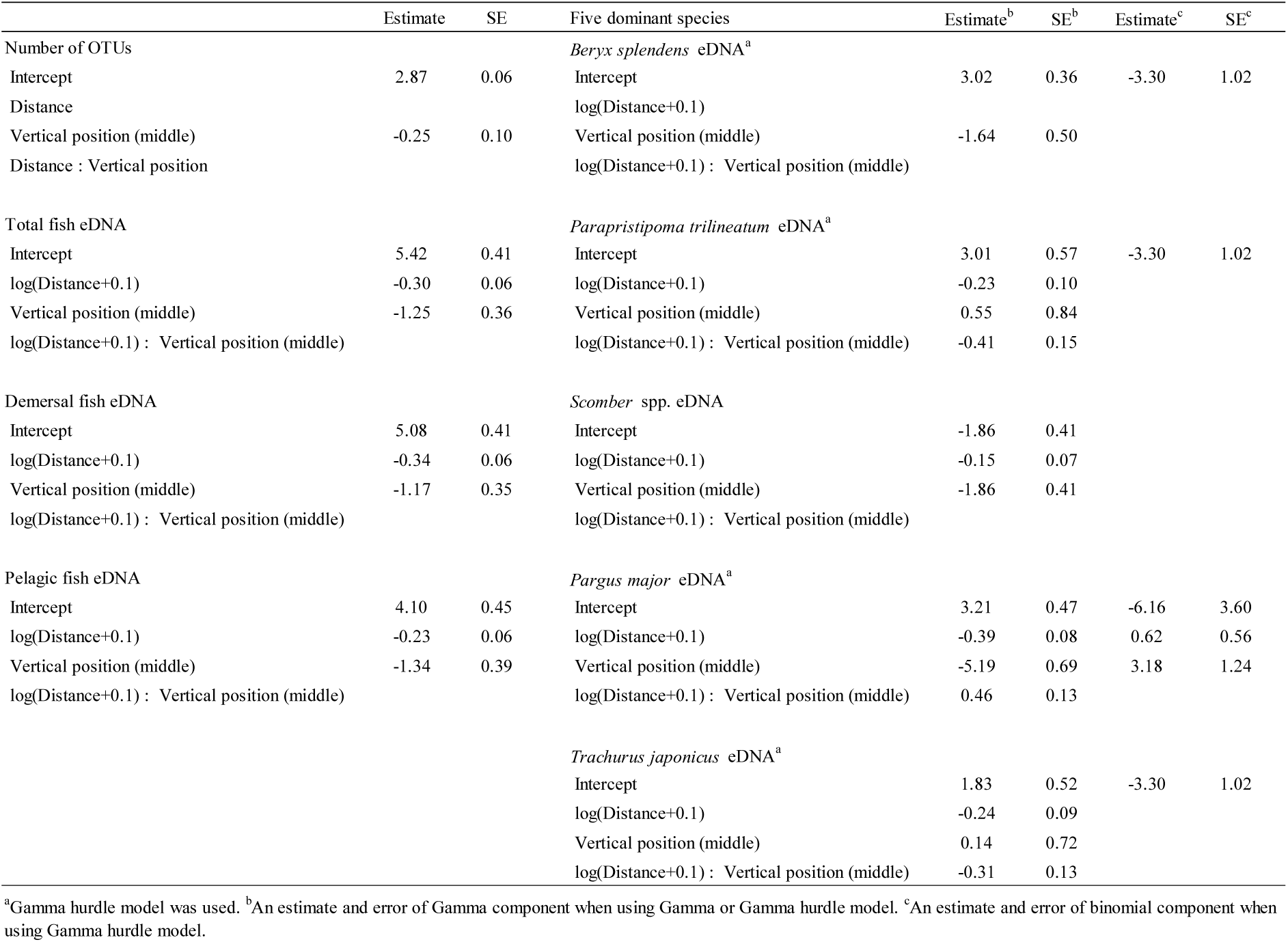
Parameters estimated from an BIC minimum model based on generalized liner modelling.

|  | Estimate | SE | Five dominant species | Estimate<sup>b</sup> | SE<sup>b</sup> | Estimate<sup>c</sup> | SE<sup>c</sup> |
| --- | --- | --- | --- | --- | --- | --- | --- |
| Number of OTUs |  |  | *Beryx splendens* eDNA<sup>a</sup> |  |  |  |  |
| Intercept | 2.87 | 0.06 | Intercept | 3.02 | 0.36 | -3.30 | 1.02 |
| Distance |  |  | log(Distance+0.1) |  |  |  |  |
| Vertical position (middle) | -0.25 | 0.10 | Vertical position (middle) | -1.64 | 0.50 |  |  |
| Distance : Vertical position |  |  | log(Distance+0.1) : Vertical position (middle) |  |  |  |  |
| Total fish eDNA |  |  | *Parapristipoma trilineatum* eDNA<sup>a</sup> |  |  |  |  |
| Intercept | 5.42 | 0.41 | Intercept | 3.01 | 0.57 | -3.30 | 1.02 |
| log(Distance+0.1) | -0.30 | 0.06 | log(Distance+0.1) | -0.23 | 0.10 |  |  |
| Vertical position (middle) | -1.25 | 0.36 | Vertical position (middle) | 0.55 | 0.84 |  |  |
| log(Distance+0.1) : Vertical position (middle) |  |  | log(Distance+0.1) : Vertical position (middle) | -0.41 | 0.15 |  |  |
| Demersal fish eDNA |  |  | *Scomber* spp. eDNA |  |  |  |  |
| Intercept | 5.08 | 0.41 | Intercept | -1.86 | 0.41 |  |  |
| log(Distance+0.1) | -0.34 | 0.06 | log(Distance+0.1) | -0.15 | 0.07 |  |  |
| Vertical position (middle) | -1.17 | 0.35 | Vertical position (middle) | -1.86 | 0.41 |  |  |
| log(Distance+0.1) : Vertical position (middle) |  |  | log(Distance+0.1) : Vertical position (middle) |  |  |  |  |
| Pelagic fish eDNA |  |  | *Pargus major* eDNA<sup>a</sup> |  |  |  |  |
| Intercept | 4.10 | 0.45 | Intercept | 3.21 | 0.47 | -6.16 | 3.60 |
| log(Distance+0.1) | -0.23 | 0.06 | log(Distance+0.1) | -0.39 | 0.08 | 0.62 | 0.56 |
| Vertical position (middle) | -1.34 | 0.39 | Vertical position (middle) | -5.19 | 0.69 | 3.18 | 1.24 |
| log(Distance+0.1) : Vertical position (middle) |  |  | log(Distance+0.1) : Vertical position (middle) | 0.46 | 0.13 |  |  |
|  |  |  | *Trachurus japonicus* eDNA<sup>a</sup> |  |  |  |  |
|  |  |  | Intercept | 1.83 | 0.52 | -3.30 | 1.02 |
|  |  |  | log(Distance+0.1) | -0.24 | 0.09 |  |  |
|  |  |  | Vertical position (middle) | 0.14 | 0.72 |  |  |
|  |  |  | log(Distance+0.1) : Vertical position (middle) | -0.31 | 0.13 |  |  |

<sup>a</sup>Gamma hurdle model was used. <sup>b</sup>An estimate and error of Gamma component when using Gamma or Gamma hurdle model. <sup>c</sup>An estimate and error of binomial component when using Gamma hurdle model.

**Table 3.** Summary results of Gamma or Gamma hurdle models (Null model: model with intercept only; Model with eco intensity: minimum BIC model which includes eco intensity as an explanatory variable) for each response variable of eDNA copy numbers based on automated model selection.

|  | Intercept<sup>b</sup> | log(eco intensity)<sup>b</sup> | Intercept<sup>c</sup> | log(eco intensity)<sup>c</sup> | BIC | R<sup>2</sup> |
| --- | --- | --- | --- | --- | --- | --- |
| Total fish eDNA |  |  |  |  |  |  |
| Null model | 4.16 |  |  |  | 325.60 | 0.00 |
| Model with eco intensity | 18.97 | 0.94 |  |  | 315.40 | 0.41 |
| Demersal fish eDNA |  |  |  |  |  |  |
| Null model | 3.80 |  |  |  | 299.20 | 0.00 |
| Model with eco intensity | 19.88 | 1.02 |  |  | 288.20 | 0.45 |
| Pelagic fish eDNA |  |  |  |  |  |  |
| Null model | 2.98 |  |  |  | 251.90 | 0.00 |
| Model with eco intensity | 15.50 | 0.79 |  |  | 244.70 | 0.33 |
| *Beryx splendens* eDNA<sup>a</sup> |  |  |  |  |  |  |
| Null model | 3.04 |  | -3.43 |  | 235.70 | 0.00 |
| Model with eco intensity | 18.84 | 1.00 | -3.43 |  | 230.70 | 0.44 |
| *Parapristipoma trilineatum* eDNA<sup>a</sup> |  |  |  |  |  |  |
| Null model | 2.66 |  | -3.43 |  | 202.20 | 0.00 |
| Model with eco intensity | 20.60 | 1.17 | -3.43 |  | 184.80 | 0.52 |
| *Scomber* spp. eDNA |  |  |  |  |  |  |
| Null model | 2.56 |  |  |  | 212.90 | 0.00 |
| Model with eco intensity | 16.32 | 0.87 |  |  | 207.60 | 0.37 |
| *Pargus major* eDNA<sup>a</sup> |  |  |  |  |  |  |
| Null model | 2.04 |  | -0.94 |  | 150.60 | 0.00 |
| Model with eco intensity | 13.99 | 0.75 | -0.94 |  | 153.60 | 0.30 |
| *Trachurus japonicus* eDNA<sup>a</sup> |  |  |  |  |  |  |
| Null model | 1.22 |  | -2.71 |  | 118.50 | 0.00 |
| Model with eco intensity | 15.62 | 0.93 | -2.71 |  | 130.70 | 0.40 |

<sup>a</sup>Gamma hurdle model was used. <sup>b</sup>An estimate and error of Gamma component when using Gamma hurdle model. <sup>c</sup>An estimate and error of binomial component when using Gamma hurdle model.

### Relationships between fish eDNA quantities and echo intensity

Based on the BIC values, echo intensity was selected as an explanatory variable for total fish, demersal fish, pelagic fish, *B. splendens*, *P. trilineatum*, *Scomber* spp., and *T. japonicus* eDNA, but not for *P. major*. The former eDNA quantities had positive relationships with echo intensity, and echo intensity had high predictive power for these eDNA quantities (Fig. 4 and Table 3).

### Fish community structure

There was dissimilarity in the NMDS in fish community structure between the AR and other surrounding stations (Fig. 5, NMDS stress = 0.096). PERMANOVA detected significant differences in community structure between the ARs (AR1 and AR2) and surrounding stations (E and W) as well as between the middle and bottom sampling layers (ARs vs surrounding stations: F = 3.040, p = 0.005; Middle vs bottom: F = 4.542, p = 0.002).

## Discussion

Field studies have begun to use an eDNA approach for monitoring fish density or diversity in coastal ecosystems^6,7,9,10^, but, as far as we know, there have been no studies using this method to assess artificial reef (AR) effects on marine fishes. We found that eDNA metabarcoding efficiently detected the fish assemblages around ARs of Tateyama Bay. MiFish metabarcoding detected 95 OTUs from 32 eDNA samples collected within 8 h on May 23, 2018 (Table S1), while 41 classifications (39 species and two classifications of closely related species) were recorded in the fish catch of the net set near the study site in May 2018 (Table S2). Of the 36 species recorded in the fish catch over a month, MiFish metabarcoding detected 58.3% of species (i.e., 21 species/36 species). This detection rate of metabarcoding relative to the conventional approach was within the range of previous MiFish metabarcoding studies (45–70%)^6,28,29^. Four dominant OTUs in terms of eDNA quantities (*P. trilineatum Scomber* spp., *P. major*, and *T. japonicus*) were also a higher proportion of fish catch of the set net (Table S1). Meanwhile, MiFish metabarcoding detected 70 fish OTUs that were not collected by the set net. This difference in the detected fish community between the MiFish metabarcoding and set net may be partly because the set net could not collect small-sized fish, and non-targeted fishes (e.g., Myctophidae and Gobiidae) were not recorded in the fish catch. These results indicated that eDNA metabarcoding approach had a high and reasonable resolution for examining a fish community in the study site.

We observed linear relationships between the number of sequence reads and the quantities of internal standard DNAs. This result suggested that the sequence reads were proportional to the copy numbers of DNA within a single sample, indicating that the conversion of sequence reads using a sample□specific regression slope would provide an estimation of DNA copy numbers in the DNA extract (i.e., estimated DNA copy number = sequence reads/sample□specific regression slope). This finding is consistent with previous studies showing that this procedure provides a reasonable estimate of copy numbers of multispecies eDNAs^7,15^. Meanwhile, the quantitative Miseq also has limitations; i.e., this method does not take account of the variation in amplification efficiency among species. Therefore, we have to caution against comparisons of DNA copy numbers between different species.

This study efficiently estimated the eDNA quantities of multiple fish species around an AR. Quantitative MiSeq sequencing found that the eDNA quantities of total, demersal, pelagic fishes, and four dominant OTUs, except *B. splendens*, were higher at AR1 than adjacent sites. In addition, all eDNA quantities, except *B. Splendens*, sharply decreased with increasing distance from the AR. In particular, those of *P. trilineatum*, *P. major*, and *T. japonicus* clearly decreased even 150 m from AR1 (e.g., W150 and E150). These results are consistent with previous studies showing a higher density of fishes in ARs than reference sites^2,30,31^. In fact, four dominant OTUs, *P. trilineatum*, *Scomber* spp., *P. major*, and *T. japonicus*, were known to appear and aggregate in ARs^2,31–33^. Only *B. splendens* was not reported to be dependent on ARs, although it is highly targeted by fisheries and usually caught by vertical longline at midnight in a deeper zone of this region (depth > 200 m, personal communication, M. Sato). A review paper of the biology of *B. splendens* indicated that the fisheries grounds of this species around the mouth of Tokyo Bay are 100–500 m in depth, but their juveniles are sometimes collected in a shallower area (<100 m)^34^. Although we cannot exclude the possibility that eDNA from adult *B. splendens* could be detected in this study, its higher eDNA quantities in the stations of 70□80 m depth (e.g., AR2, AR1 and E150) would be likely to reflect the presence of its juveniles. In future studies, combining eDNA analysis with other methods such as observation by underwater drones or GoPro, which can determine their body length, would clarify the details of ontogenetic habitat shift for *B. splendens*.

We expected the number of OTUs would be higher at ARs than the surrounding stations, but did not observe such a pattern. In the following year’s survey, we found a larger number of fishes around AR1 than surrounding stations in a GoPro video (personal observation, N. Inoue, M. Sato, N, Furuichi), which contradicted our eDNA results. We suspect that higher eDNA concentrations of dominant OTUs at ARs consumed a large portion of the sequence reads, and species with low eDNA concentrations were not detected by MiFish metabarcoding in this study.

MiFish metabarcoding also showed different fish species composition between the AR and surrounding stations based on NMDS and PERMANOVA. The eDNA quantities of fishes, including dominant OTUs (pelagic fishes: *Scomber* spp., *Etrumeus teres*, *Sardinops melanostictus*; demersal fishes: *B. splendens*, *P. trilineatum*, *P. major*, *Sardinops melanostictus*, *Sacura margaritacea*, *Thamnaconus modestus*) were more abundant in ARs, while those of some demersal fishes such as flathead (*Platycephalus* sp.2), Asian sheephead wrasse (*Semicossyphus reticulatus*), barred knifejaw (*Oplegnathus fasciatus*), and seabream (*Dentex tumifrons* or *D. cardinalis*) were higher in the surrounding stations. Different fish responses to AR structures may result in differences in the fish community compositions between the AR and surrounding stations.

Previous studies have found positive relationships between the density of target species and eDNA concentrations in semi-closed marine systems like estuaries^9,13^; our results found such associations in an open marine system. We found eDNA quantities of total, demersal, pelagic fishes, and four dominant species, except *P. major*, positively correlated with the echo intensity measured by echo sounder. Our study site faces toward the open sea (i.e., Pacific ocean), and the flow was stronger than in a previous study (5.0–25.0 cm/s in our study and 0.4–13.5 cm/s in a semi-closed bay at Maizuru, Japan)^35^. Therefore, the flow is likely to transport more eDNAs from sampling stations in this study, but we could also obtain a strongly localized eDNA signal at sampling stations. One exception is the eDNA quantity of *P. major*, which did not correlate with echo intensity. This finding was probably due to elimination of the echo signals around the sea bottom, where this species is mainly found^18^. Overall, our result indicates that eDNA quantities reflect spatial distributions of marine fish in an open coastal environment.

We found the eDNA quantities and species number were higher in the bottom sampling layers than in the middle layers. Multivariate analyses, including NMDS and PERMANOVA, also showed different fish compositions between the bottom and middle layers. Previous eDNA studies have also indicated variations in eDNA quantities, the number of species, and community compositions between different sampling depths^6,9,36,37^. Some of these studies using eDNA methods suggested that water column stratification and the different species’ responses to it cause the observed different composition between sampling depths^36,37^. Murakami et al.^35^ also discussed that the vertical distribution of eDNA depends on both the depth of the source organisms and the behavior of eDNA, with the latter dependent on its chemical condition and physical factors in the environment. Although water column stratification between the middle and bottom layers was not observed in our study (Fig. S3), we consider that the preference of detected fish species for bottom habitats or the behavior of eDNA contributed to the higher eDNA quantities and OTU number in the bottom layer in this study.

Overall, our results demonstrate the effectiveness of quantitative MiSeq sequencing to assess AR effects on multiple fish species. We found higher quantities of fish eDNA at AR stations than at surrounding stations and positive correlations between these quantities and eco intensity, which indicates a highly localized eDNA signal in an oceanographically dynamic environment. This spatial resolution indicates that quantitative MiSeq sequencing can improve techniques to monitor the distribution of multiple fish species. Another method, called ‘quantitative sequencing (qSeq).’ has also been developed for simultaneous quantifications of eDNA concentrations of multiple species^38^. This method adopts a stochastic labeling approach with random-tag sequences and can also quantify eDNA concentrations of fish species using the MiFish primer set^39^. The quantitative metabarcoding methods (i.e., the Miseq sequencing and qSeq) can complement conventional methods, especially in an environment with areas where the water is too deep to adopt underwater visual census (>40m depth) or in no-take marine reserves where exploiting fish is prohibited.

## Data availability

DDBJ accession numbers of the DNA sequences analyzed in the present study are DRA011864 (Submission ID), PRJDB11389 (BioProject ID) and SAMD00298249–SAMD00298281 (BioSample ID).

## Acknowledgements

We are grateful to the staff of the research vessel *Takamaru* for their support in the field, Tomoe Ishibashi of Fisheries Technology Institute, and Tomohiro Shirako of IDEA consultants, Inc. for support in molecular experiments. We also thank Yutaro Kawano of Mie University and Fumie Yamaguchi of Fisheries Technology Institute for providing fish photos. This work was supported by the research grants from the National Research Institute of Fisheries Engineering (NRIFE) in 2018□2019 fiscal year.

## Author Contributions

M. S., R. N., and N. F. designed the study and conducted field surveys. T. I. and N. I. analyzed the data of echo sounder survey. M. S. and N. I. performed molecular experiments, and M. U. prepared the internal standards for eDNA quantification. M. S. and N. I. analyzed fish community dataset extracted from results of MiSeq sequencing. M. S. wrote the first draft of the manuscript and other authors commented on the manuscript.

## Additional information

### Supplementary information

accompanies this paper.

