## Supplementary materials for "Quantitative assessment of multiple fish species around artificial reefs using environmental DNA metabarcoding"

**Title**

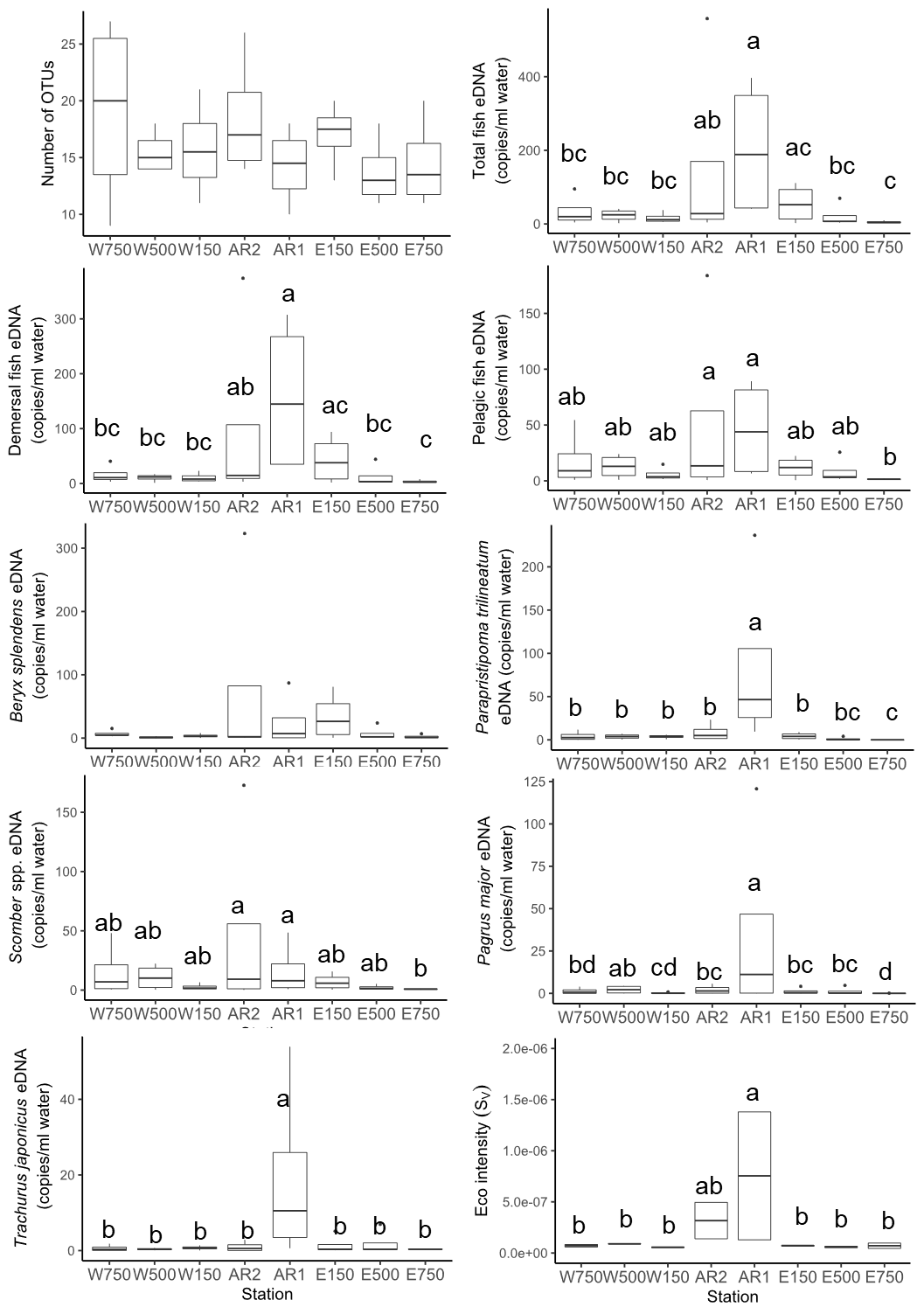

Figure S1 Box plots indicate number of OTUs, eDNA copies of total fish, demersal fish, pelagic fish, Beryx splendens, Parapristipoma trilineatum, Scomber spp., Pagrus major, and Trachurus japonicus, and eco intensity at each station. Different lowercase letters represent statistically significant differences (P < 0.05; Tukey tests).

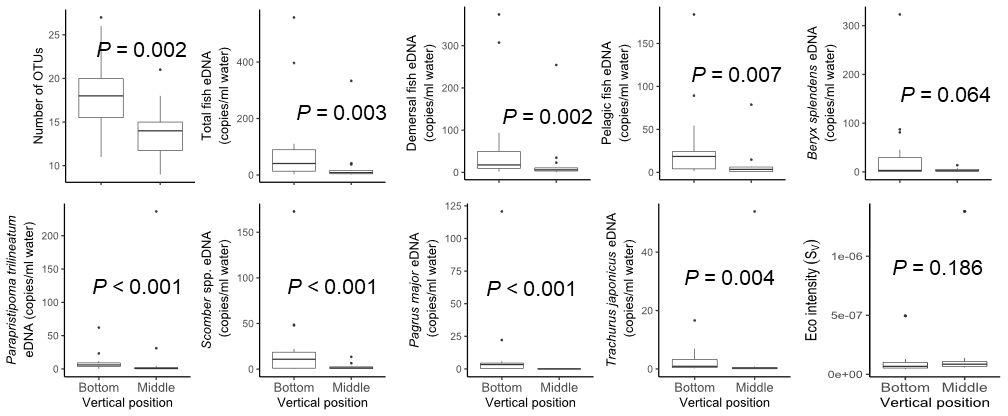

Figure S2 Box plots indicate number of OTUs, eDNA copies of total fish, demersal fish, pelagic fish, Beryx splendens, Parapristipoma trilineatum, Scomber spp., Pagrus major, and Trachurus japonicus, and eco intensity from bottom and middle samples. P-values indicate statistical significance for the differences between bottom and middle samples.

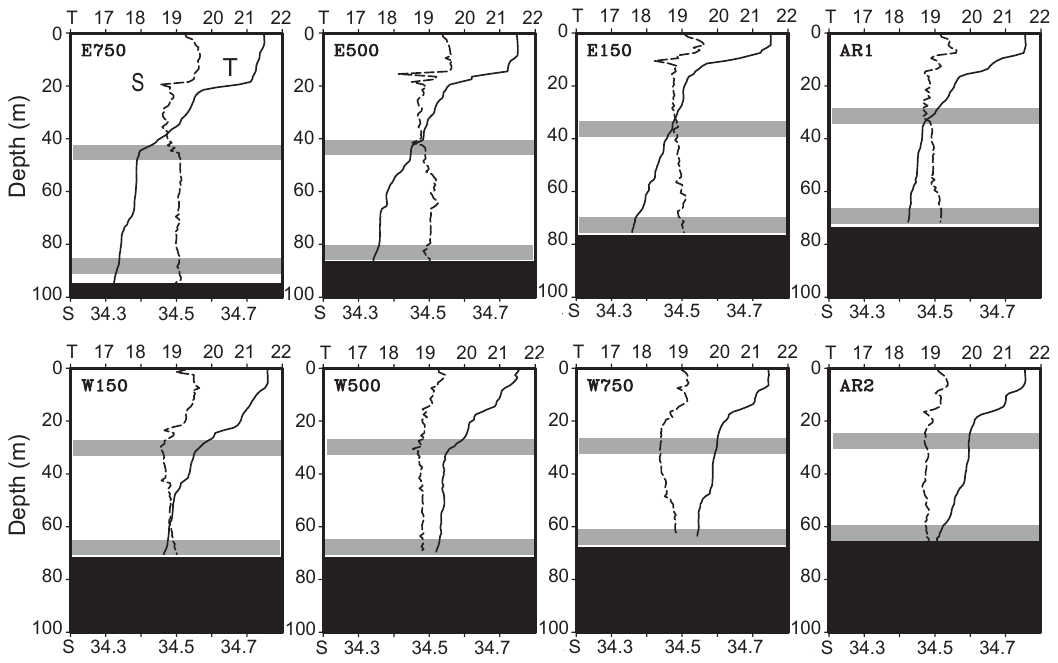

Figure S3 Vertical profiles of seawater temperature (T, °C, solid line) and salinity (S, dashed line). The gray and black shading indicate the depth range of water sampling and bottom topography, respectively.

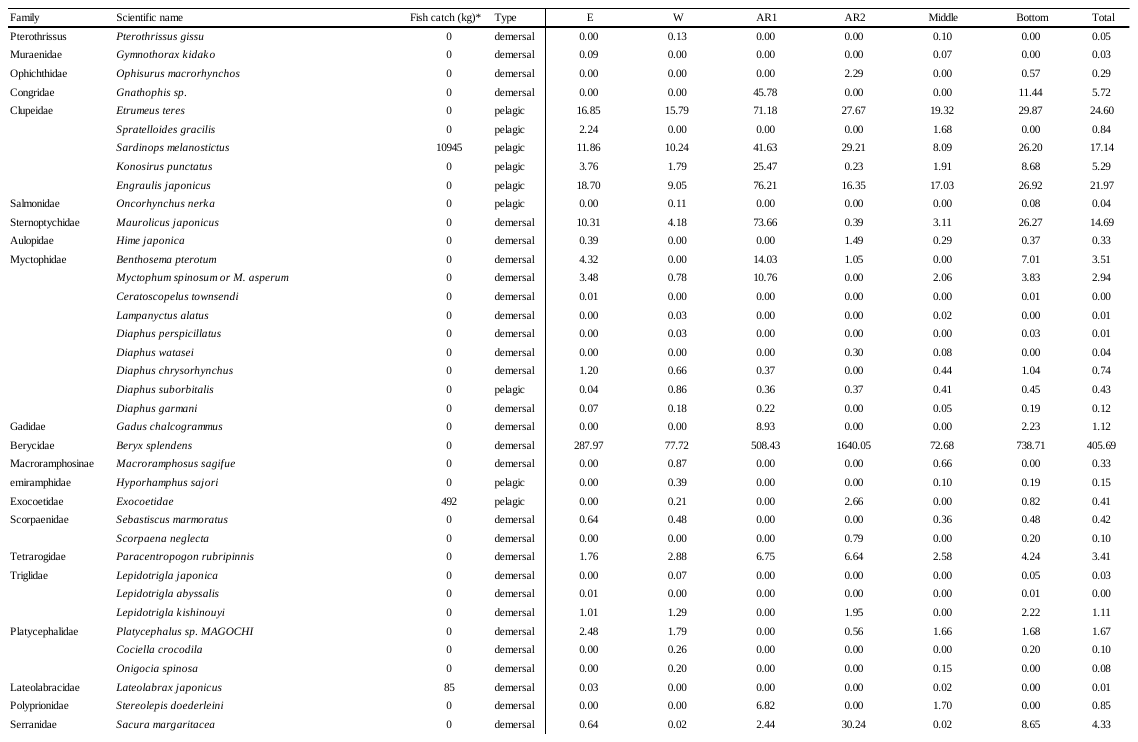
Table S1. Average eDNA copy numbers of each fish species in study stations and vertical positions at Tateyama Bay. *Fish catch (kg) by a set net near study site in May 2018.

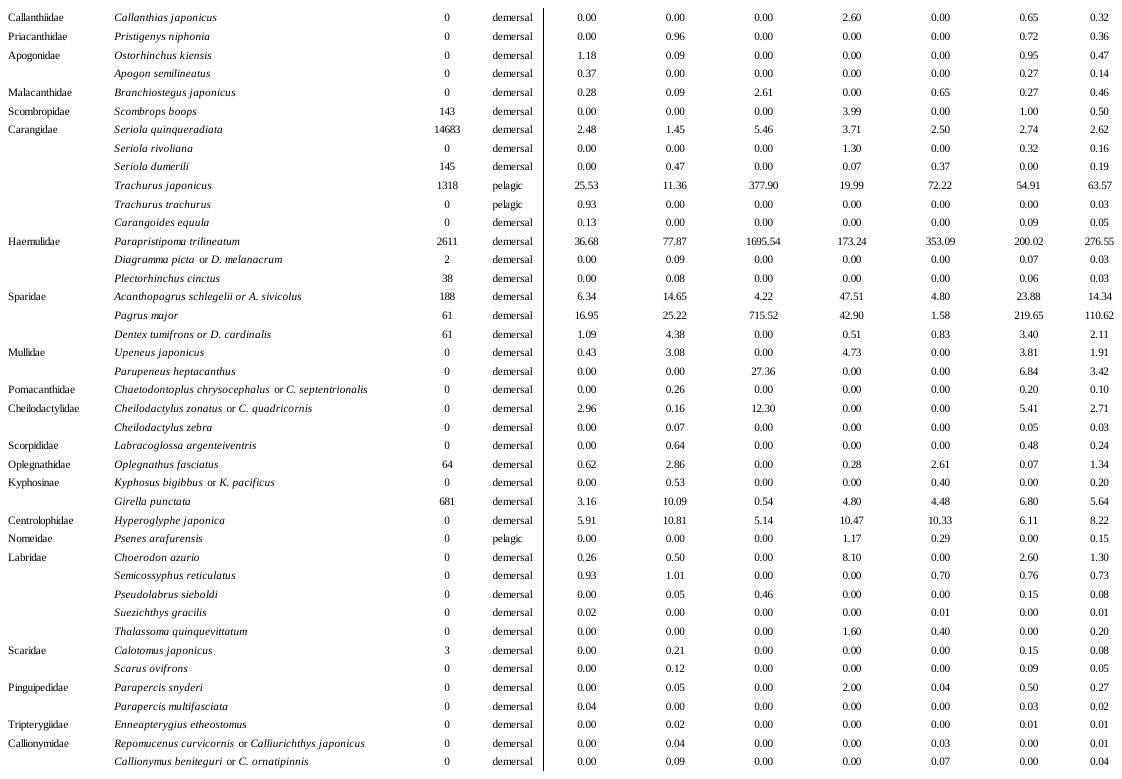

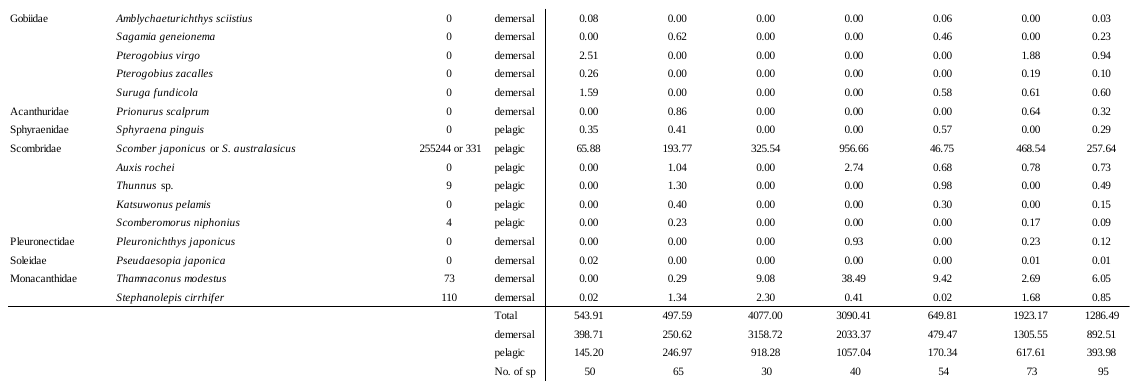

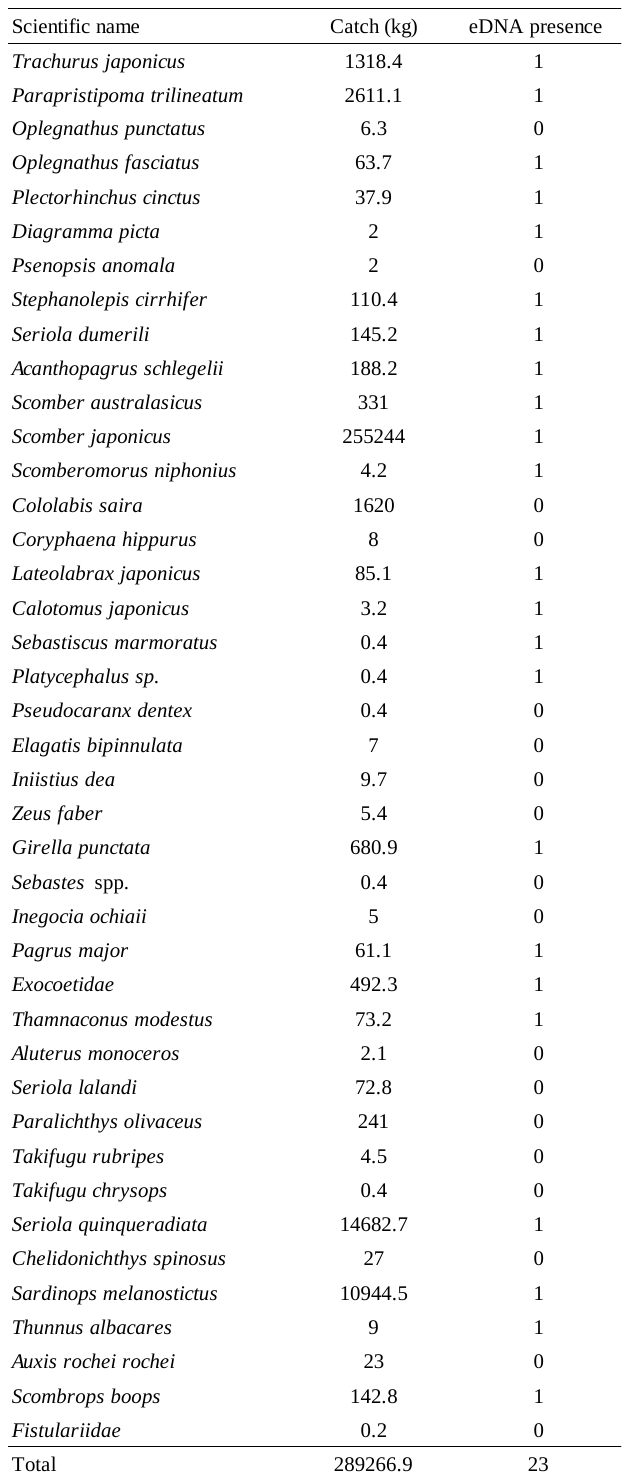
Table S2. Species list of fish catch (kg) by a set net near study site in May 2018.

Table S3. Results of likelihood ratio tests examining variation in number of OTUs, eDNA copies of fishes, and eco intensity among sampling stations as well as those between vertical sampling positions.

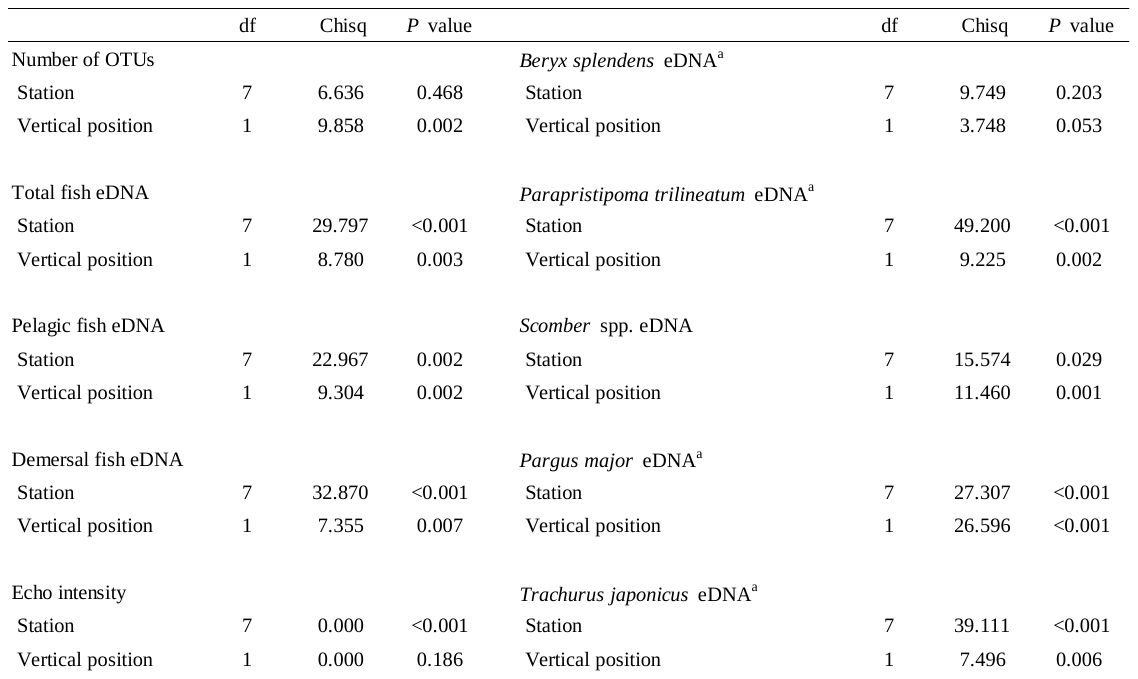
